# Differential transpiration between pods and leaves during stress combination in soybean

**DOI:** 10.1101/2022.12.20.521196

**Authors:** Ranjita Sinha, Benjamin Shostak, Sai Preethi Induri, Sidharth Sen, Sara I. Zandalinas, Trupti Joshi, Felix B. Fritschi, Ron Mittler

## Abstract

Climate change is causing an increase in the frequency and intensity of droughts, heat waves, and their combinations, diminishing agricultural productivity and destabilizing societies worldwide. We recently reported that during a combination of water deficit (WD) and heat stress (HS) stomata on leaves of soybean plants are closed, while stomata on flowers are open. This unique stomatal response was accompanied by differential transpiration (higher in flowers, while lower in leaves) that cooled flowers during a combination of WD+HS. Here we reveal that developing pods of soybean plants subjected to a combination of WD+HS use a similar acclimation strategy of differential transpiration to reduce internal pod temperature by about 4°C. We further show that enhanced expression of transcripts involved in abscisic acid degradation accompanies this response, and that preventing pod transpiration by sealing stomata causes a significant increase in internal pod temperature. Using an RNA-Seq analysis of pods developing on plants subjected to WD+HS, we also show that the response of pods to WD, HS, or WD+HS is distinct from that of leaves or flowers. Interestingly, we report that although flower, pod and seed numbers per plant are decreased under conditions of WD+HS, seed mass of plants subjected to WD+HS is larger than that of plants subjected to HS, and number of seeds with suppressed/aborted development is lower in WD+HS compared to HS. Taken together our findings reveal that differential transpiration occurs in pods of soybean plants subjected to WD+HS and that this process limits heat-induced damage to seed production.

**One sentence summary:** Differential transpiration between pods and leaves of soybean plants subjected to a combination of water deficit and heat stress buffers internal pod temperature.

## INTRODUCTION

Global warming and climate change are gradually altering our environment, causing an increase in the frequency and intensity of devastating weather events such as floodings, extended droughts, and heat waves (Alizadeh et al., 2020; Overpeck and Udall, 2020; Zhai et al., 2021). These events negatively impact agricultural production and destabilize different societies worldwide (Lobell et al., 2011; Bailey-Serres et al., 2019; Zandalinas et al., 2021). Of particular concern to agricultural productivity is the increase in the frequency of drought and heat wave combination episodes, in recent years (Mazdiyasni and AghaKouchak, 2015; Alizadeh et al., 2020; Rivero et al., 2021). Historically, episodes of drought and heat stress combination had a catastrophic impact on agricultural production (*e*.*g*., the drought and heat wave episodes of 1980 and 1988 in the US that resulted in losses to agriculture estimated at $33 and 44 billion, respectively; Mittler, 2006; https://www.ncdc.noaa.gov/billions/), and their increased frequency requires special attention. Multiple studies have now shown that the molecular, physiological, and metabolic response of plants subjected to water deficit (WD), or heat stress (HS) is different from that induced in plants during a combination of WD and HS (WD+HS), and could involve conflicting pathways and/or responses (Mittler, 2006; Zandalinas et al., 2020b; Zandalinas et al., 2021; Sinha et al., 2022; Zandalinas and Mittler, 2022; Mittler et al., 2022). Moreover, it was found that when droughts and heat waves co-occur during the reproductive growth phase of crops, their impact is significantly higher than when they co-occur during vegetative growth (Mahrookashani et al., 2017; Lawas et al., 2018; Liu et al., 2020; Cohen et al., 2021b; Sinha et al., 2021).

Among the many conflicting responses of plants to WD and HS is the regulation of stomatal aperture. During WD stomata close to prevent water loss, but during HS stomata open to cool the leaf by transpiration (Nilson and Assmann, 2007; Lawson and Matthews, 2020; Xie et al., 2022). During a combination of WD and HS, stomata on leaves of many plants remain however closed and leaf temperature increases to levels that are even higher than that of HS alone; because the plant cannot cool its leaves by transpiration (Mittler, 2006; Sinha et al., 2022; Mittler and Zandalinas, 2022). We recently reported that during a combination of WD+HS, stomata on leaves of soybean plants are closed, while stomata on flowers of soybean (sepals) are open (Sinha et al., 2022). This differential regulation of stomatal aperture is accompanied by differential transpiration (higher in flowers, while lower in leaves) that allowed soybean plants subjected to WD+HS to cool their flowers and limit heat-induced damages to reproductive organs (Sinha et al., 2022). We termed this acclimation strategy ‘Differential transpiration’ and identified enhanced rates of abscisic acid (ABA) degradation in flowers from plants subjected to WD+HS, or just HS, as playing a key role in this response, allowing stomata on flowers to remain open during conditions of WD+HS (or HS). A similar response was not observed in plants subjected WD alone and the stomata on flowers and leaves from these plants remained closed (Sinha et al., 2022).

While reproductive organ differentiation in flowers, as well as the different processes involved in in plant fertilization, are highly sensitive to heat stress (Gray and Brady, 2016; Santiago and Sharkey, 2019; Chaturvedi et al., 2021; Sze et al., 2021), so are processes that occur in pods following successful fertilization (Siebers et al., 2015; Sehgal et al., 2018; Djanaguiraman et al., 2019, 2020). For example, the number of seeds per pod, the size of seeds, and the overall process of seed filling, were shown to be reduced by HS (Siebers et al., 2015; Sehgal et al., 2018; Djanaguiraman et al., 2019, 2020). Because differential transpiration was found to play an important role in the cooling of soybean flowers (Sinha et al., 2022), we hypothesized that the same mechanism could also be involved in limiting heat-induced damage to developing pods and seeds during conditions of WD+HS. Here we reveal that developing pods of soybean plants subjected to a combination of WD+HS use differential transpiration to buffer their internal temperature. We further show that this process is associated with enhanced expression of transcripts involved in ABA degradation, and that preventing it by sealing stomata causes a significant increase in internal pod temperature. Using an RNA-Seq analysis of pods from plants subjected to control (CT), WD, HS, or WD+HS we further show that the response of pods to WD, HS, or WD+HS is distinct from that of leaves or flowers. Interestingly, we report that although the number of flowers, pods and seeds per plant are decreased under conditions of WD+HS, seed mass of plants subjected to WD+HS is larger than that of plants subjected to HS. Moreover, the number of seeds with suppressed/aborted development in pods from plants subjected to WD+HS was lower than that of plants subjected to HS. Taken together, our findings reveal that differential transpiration occurs in pods of plants subjected to WD+HS and that this process buffers internal pod temperature and protects seed development under conditions of WD+HS combination.

## RESULTS

### Characterization of pods developed on soybean plants subjected to WD, HS, or a combination of WD+HS

To study the effects of WD+HS on developing pods, we grew soybean plants (*Glycine max* cv Magellan) in four identical growth chambers under controlled growth conditions until plants began to flower (Cohen et al., 2021a; Sinha et al., 2022). At the beginning of flowering (R1 stage; Fehr et al., 1971) we randomized the plants into conditions of WD, HS, WD+HS, or CT in the four growth chambers (Cohen et al., 2021; Sinha et al., 2022) and maintained these conditions until the end of the experiments (6 weeks). At 20 days following the initiation of stress treatments we started sampling and analyzing pods from all chambers. This design allowed us to study pods that developed on plants under the different stress conditions. As shown in Figure 1A, all pods used for our physiological and molecular studies were at a length of about 3 cm and contained developing seeds. Thermocouple thermometer probe measurements of internal pod temperature revealed that pods from plants subjected to WD had a higher internal temperature compared to CT (Figure 1B). As expected, the internal pod temperature of plants subjected to HS was higher than that of CT or WD conditions (Figure 1B). Interestingly, the internal temperature of pods from plants subjected to WD+HS was not significantly different than that of plants subjected to HS (Figure 1B). As shown in Figure 1C, the water potential of pods from plants subjected to WD+HS was lower than that of pods from plants subjected to CT or WD conditions, while the water potential of pods from plants subjected to HS was at an intermediate level between WD and WD+HS.

**Figure 1.**
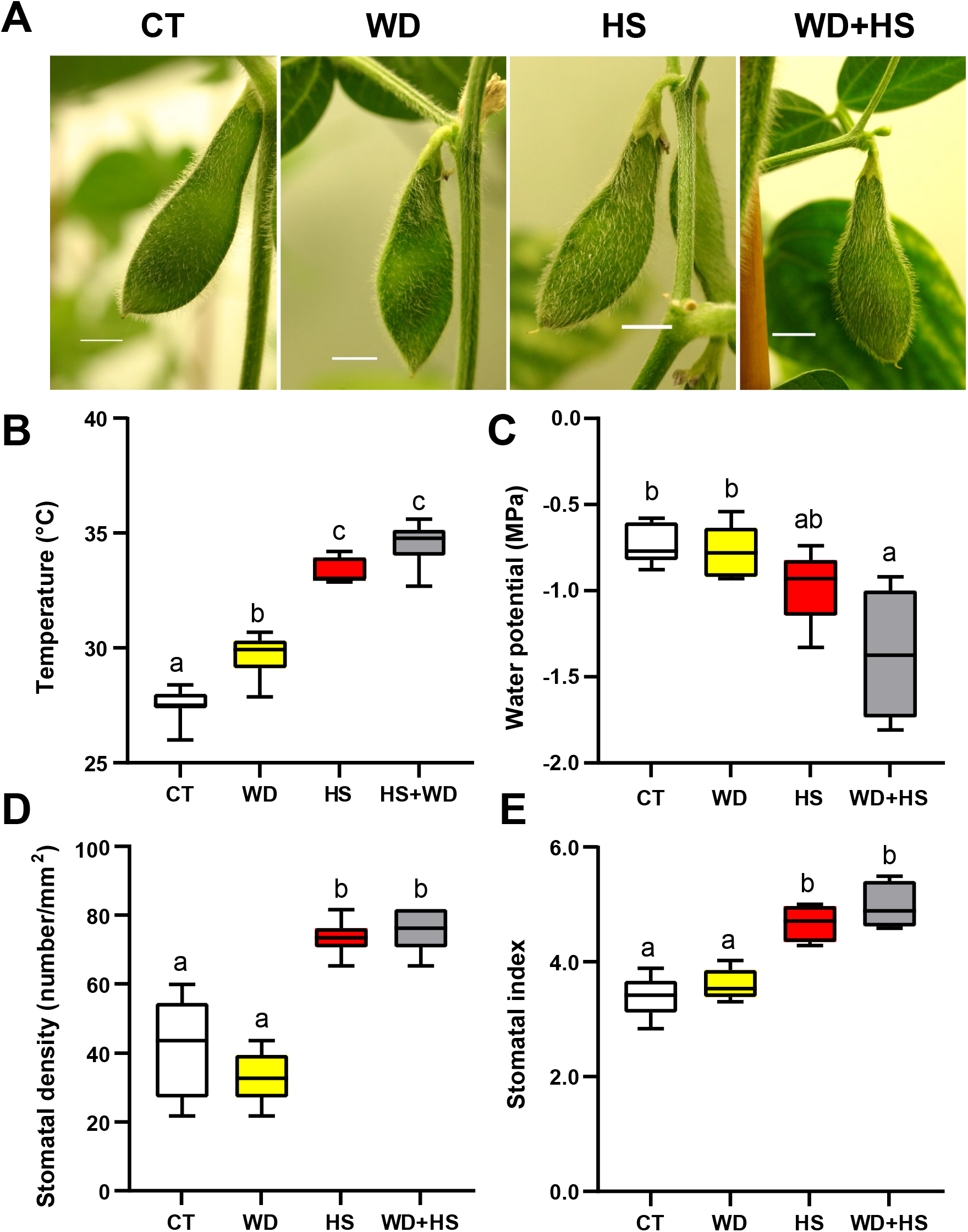
Inner temperature, water potential, and stomatal density and index of pods from plants grown under a combination of water deficit and heat stress conditions. **(A)** Representative images of pods from soybean plants developing under control (CT), water deficit (WD), heat stress (HS) and a combination of WD+HS conditions. Bar is 5 mm. **(B-E)** Inner temperature (B), water potential (C), stomatal density (D), and stomatal index (E) of pods from soybean plants subjected to CT, HS, WD, or WD+HS conditions. All experiments were conducted with 3 biological repeats, each with at least 15 plants as technical repeats. Results are shown as box-and-whisker plots with borders corresponding to the 25^th^ and 75^th^ percentiles of the data. Different letters denote significance at P < 0.05 (ANOVA followed by a Tukey’s post hoc test). Abbreviations: CT, control; HS, heat stress; MPa, mega pascal; WD, water deficit.

We previously reported that the stomatal density of flowers (sepals) developing on plants subjected to HS or WD+HS was higher compared to that of flowers from plants subjected to WD or CT conditions (Sinha et al., 2022). This observation correlated with higher rates of transpiration in flowers from plants subjected to HS or WD+HS (Sinha et al., 2022). To test whether pods developing on plants subjected to HS or WD+HS would also contain a higher number of stomata, we measured stomatal density and index of these pods. Indeed, the stomatal density (Figure 1D) and index (Figure 1E) of pods developing on plants subjected to HS or WD+HS were higher compared to that of pods developing on plants subjected to CT or WD conditions.

### Transpiration and stomatal conductance of pods from plants subjected to WD+HS

Transpiration and stomatal conductance of pods and leaves from plants subjected to WD+HS were measured between 12:00 and 13:00 hours. Each time, the transpiration and stomatal conductance of pods and leaves from the same plants were measured and compared. While the transpiration of leaves from plants subjected to WD+HS was suppressed, the transpiration of pods from the same plants was not (Figure 2A). In contrast, the transpiration of leaves and pods from plants subjected to HS was not suppressed, and the transpiration of leaves from plants subjected to WD was (Figure 2A). Not surprisingly, stomatal conductance of pods and leaves from the different treatments corresponded to the transpiration rates measured in these plants (Figure 2B). The findings presented in Figures 1 and 2 reveal that, like flowers from plants subjected to WD+HS (Sinha et al., 2022), pods from plants subjected to WD+HS also continue to transpire potentially to control their internal temperature. Differential transpiration, that was discovered between flowers (sepals) and leaves during conditions of WD+HS (Sinha et al., 2022), therefore also occurs between pods and leaves under conditions of WD+HS (Figures 1 and 2).

**Figure 2.**
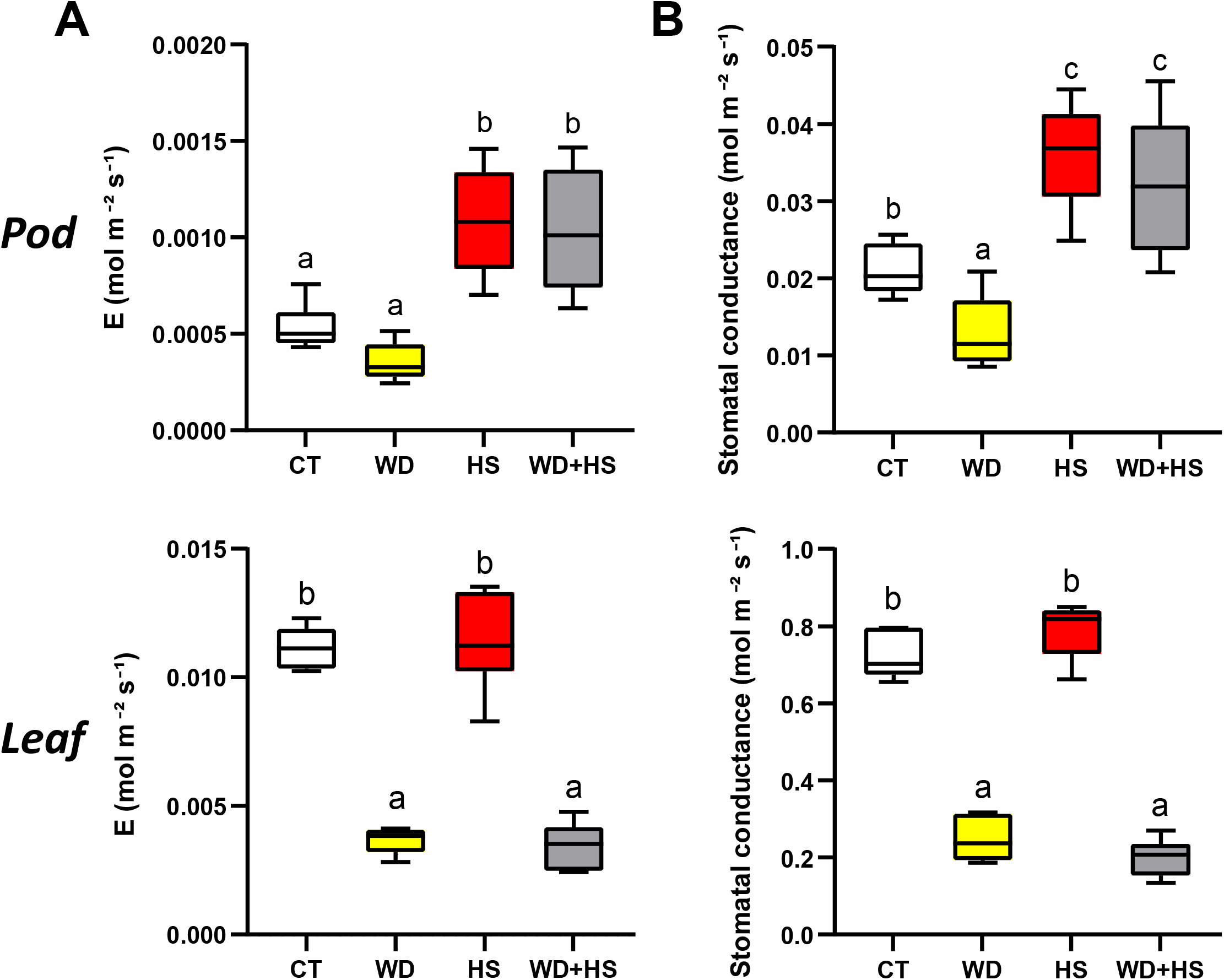
Transpiration and stomatal conductance of pods from plants grown under a combination of water deficit and heat stress conditions. **(A)** Transpiration of pods (top) and leaves (bottom) from soybean plants developing under control (CT), water deficit (WD), heat stress (HS) and a combination of WD+HS conditions. **(B)** Stomatal conductance of pods (top) and leaves (bottom) from soybean plants developing under CT, WD, HS, and WD+HS conditions. All experiments were conducted with 3 biological repeats, each with at least 15 plants as technical repeats. Results are shown as box-and-whisker plots with borders corresponding to the 25^th^ and 75^th^ percentiles of the data. Different letters denote significance at P < 0.05 (ANOVA followed by a Tukey’s post hoc test). Abbreviations: CT, control; E, transpiration; HS, heat stress; WD, water deficit.

### RNA-Seq analysis of pods from plants subjected to WD+HS

We previously studied the transcriptomic response of soybean leaves and flowers to CT, WD, HS, and WD+HS conditions, using the same growth chambers, conditions, and protocols described in this work (Cohen et al., 2021a; Sinha et al., 2022). These studies revealed that the transcriptomic response of soybean leaves and flowers to WD+HS is different from that to WD or HS (Cohen et al., 2021a; Sinha et al., 2022). To test whether pods would also display such a unique transcriptomic response to the stress combination, we conducted RNA-Seq analysis of pods obtained from plants subjected to CT, WD, HS, or WD+HS conditions (Figures 1 and 2). As shown in Figure 3A, the transcriptomic response of pods to a combination of WD+HS was extensive with over 11,000 transcripts altered in their abundance (Supplemental Tables 1-6). Compared to this response, the response of pods to WD or HS was much lower with 359 and 2402 transcripts altered in their abundance, respectively. Interestingly, the expression of only 118 transcripts was commonly altered in pods in response to all three stress treatments, demonstrating a low similarity between the transcriptomic response of pods to WD, HS, and WD+HS. Transcripts specifically expressed during the WD+HS combination in pods (Figure 3A; 9,959) were enriched in mitochondrial-related processes, ubiquitin-dependent protein degradation, cell wall, lipid, and ion transport functions (Figure 3B; Supplemental Table 7).

**Figure 3.**
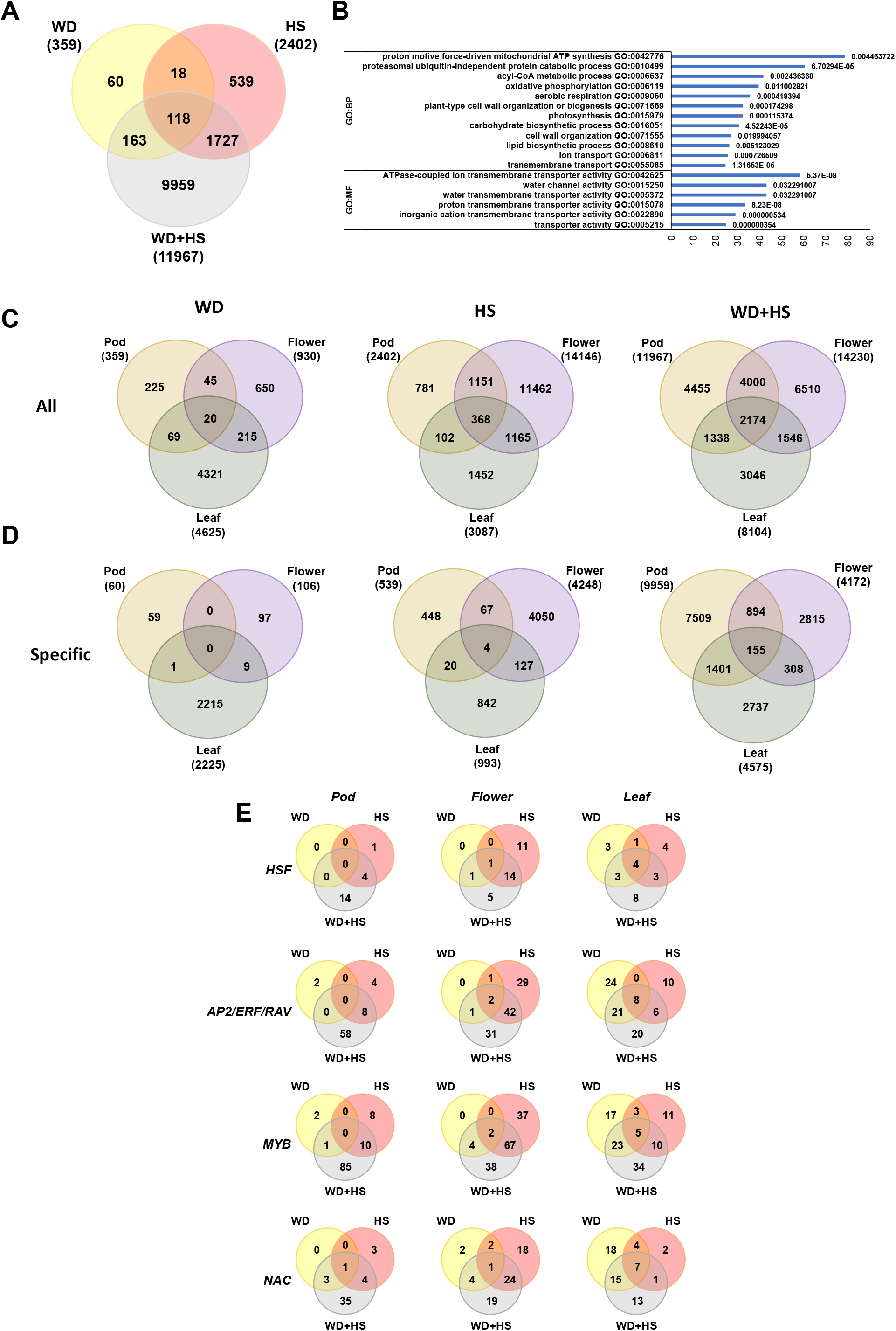
RNA-Seq analysis of pods from soybean plants subjected to a combination of water deficit and heat stress. **(A)** Venn diagram showing the overlap between transcripts with significantly altered expression (up or down regulated) in pods from plants grown under control (CT), water deficit (WD), heat stress (HS) and a combination of WD+HS conditions. **(B)** Representative GO enrichment analysis results of transcripts unique to WD+HS in pods (9959; See Supplementary Table S7 for a complete list). **(C)** Venn diagrams showing the overlap between transcripts with significantly altered expression (up or down regulated) in pods, flowers and leaves from plants grown under CT, WD, HS and WD+HS conditions. **(D)** same as in (C) but for transcripts unique to WD, HS, and WD+HS combination. **(E)** Venn diagrams showing the overlap in expression of selected transcription factor (TF) families in pods, flowers, and leaves from plants grown under CT, WD, HS and WD+HS conditions. Additional Venn diagrams are sown in Supplementary Figure 1. All transcripts shown are significant at P < 0.05 (negative binomial Wald test followed by Benjamini–Hochberg correction). Abbreviations: AP2/ERF/RAV, APETALA2/ ethylene response factor/ related to ABI3 and VP1; CT, control; GO, gene ontology; HS, heat stress; HSF, heat shock transcription factor; MYB, v-Myb myeloblastosis viral oncogene homolog; NAC, NAM, ATAF and CUC TF; TF, transcription factor; WD, water deficit.

To determine how similar or different were leaves, flowers, and pods in their transcriptomic responses to WD, HS, or WD+HS, we generated Venn diagrams comparing the response of each tissue to each of the three different stress conditions (Figure 3C). Interestingly, as shown in Figure 3C (left Venn diagrams), compared to the transcriptomic response of leaves to WD (Cohen et al., 2021a; Sinha et al., 2022; 4,624 transcripts), the transcriptomic response of pods and flowers to WD was less extensive, with only 359 and 930 transcripts, respectively. In addition, compared to the transcriptomic response of flowers to HS that was extensive (Sinha et al., 2022; 14,146 transcripts), the transcriptomics response of pods and leaves to HS was much more muted with 2,402 and 3,087 transcripts, respectively (Figure 3C; middle Venn diagrams). In response to WD+HS, however, the transcriptomic response of all three tissues was extensive with 14,230, 11,967, and 8,104 transcripts altered in expression, in flowers, pods and leaves, respectively (Figure 3C; right Venn diagrams; Cohen et al., 2021a; Sinha et al., 2022). Although the three different responses to WD+HS shared over 2,000 transcripts in common, each tissue displayed a distinct transcriptomic response to WD+HS that included over 4,000, 6,500, and 3,000 transcripts, unique to pods flowers, and leaves, respectively (Figure 3C; right Venn diagrams).

When the differentially expressed pod transcripts, unique to WD, HS or WD+HS (60, 539, 9,959 transcripts, respectively; Figure 3A) were compared to the differentially expressed leaves and flower transcripts, unique to the same conditions (Cohen et al., 2021a; Sinha et al., 2022), the overlaps in transcriptomic responses were even lower, demonstrating that the distinct responses of pods to each of the different stress conditions shared limited similarity with the responses observed in other tissues (Figure 3D; Supplemental Tables 8-16; Cohen et al., 2021a; Sinha et al., 2022). Gene ontology (GO) annotation of the transcripts altered in each tissue under CT, WD, HS, and WD+HS conditions (Supplemental Tables 7 and 17-20) further supports these findings.

A comparison between the expression pattern of transcripts encoding different transcription factor (TF) families (Zandalinas et al., 2020a; Zhang et al., 2021; Mittler et al., 2022) in leaf, flower, and pod tissues during WD, HS or WD+HS further revealed that the response of each tissue to WD+HS was distinct and accompanied by changes in the expression of many TFs (Figure 3E; Supplemental Figure S1; Supplemental Tables 21-28). Interestingly, while the response of flowers and leaves to HS was relatively extensive, the response of pods to HS involved fewer TFs. In addition, while the response of leaves to WD was extensive, the responses of flowers and pods were not (Figure 3E; Supplemental Figure S1; Supplemental Tables 21-28). This finding is intriguing since it suggests that compared to leaves, pods (and flowers) respond differently to the effects of WD, while compared to flowers, pods respond differently to the effects of HS. Nevertheless, once WD and HS are combined (WD+HS), the response of all plant tissues was extensive (Figure 3).

### Expression of transcripts encoding ABA biosynthesis and degradation enzymes in pods from plants subjected to WD, HS or WD+HS

We previously reported that flowers from plants subjected to HS or a combination of WD+HS (but not WD) displayed a higher abundance of transcripts encoding the ABA degradation enzyme CYP707A (ABA 8’-hydroxylase), were less sensitive to external ABA application (*i*.*e*., displayed stomatal closure only in response to the application of higher ABA concentrations, compared to CT), and contained higher levels of the ABA degradation byproduct dihydrophaseic acid (DPA; Sinha et al., 2022). These findings suggested that at least part of the differential transpiration phenotype between leaves and flowers, displayed by plants subjected to a combination of WD+HS, is mediated by enhanced rates of ABA degradation that occur in flowers under these conditions (Sinha et al., 2022). To test whether the differential transpiration between pods and leaves, displayed in soybean plants subjected to a combination of WD+HS (Figure 2), is also associated with a similar mechanism, we compared the expression of ABA metabolizing enzymes between pods, leaves, and flowers during the different stress treatments, as well as determined the sensitivity of pods to external application of ABA. As shown in Figure 4A, in contrast to leaves and flowers, the expression of all ABA biosynthesis genes was not elevated in pods in response to WD. In addition, while the expression of biosynthesis genes encoding ABA1 (Zeaxanthin epoxidase) and NCED (9-cis-Epoxycarotenoid dioxygenase) was elevated in response to WD+HS, the expression of ABA2 (Xanthoxin dehydrogenase 2) and AAO3 (Aldehyde oxidase 3) was not. In agreement with our previous findings with flowers (Sinha et al., 2022), the expression of CYP707A (ABA 8’-hydroxylase), involved in ABA degradation, was elevated in response to HS and WD+HS (but also in response to WD) in pods.

**Figure 4.**
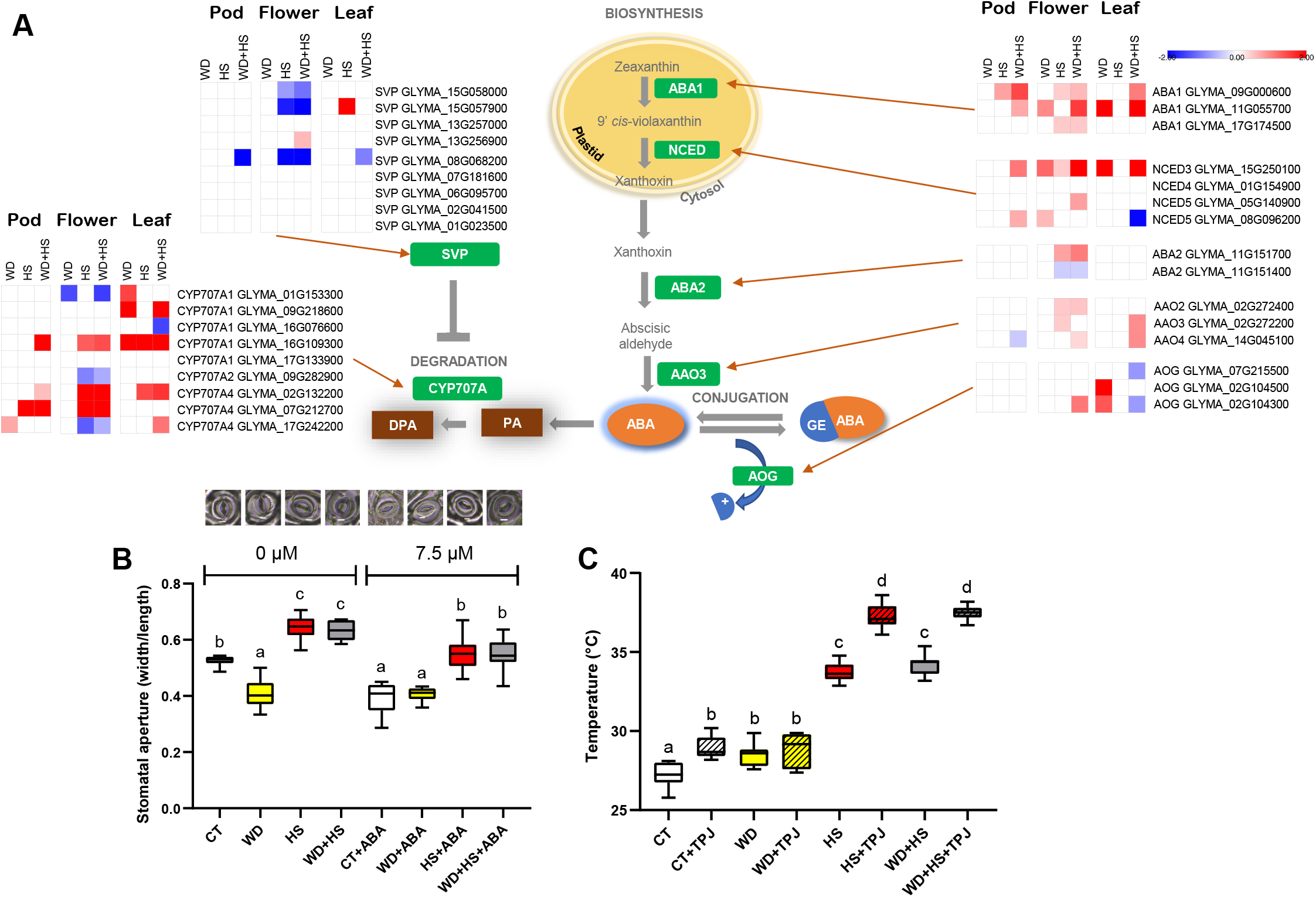
Expression of transcripts involved in ABA metabolism, reduced sensitivity to ABA, and effects of sealing stomata on internal pod temperature, in pods from plants subjected to a combination of water deficit and heat stress conditions. **(A)** Heat maps and a pathway showing the expression of transcripts involved in ABA biosynthesis and degradation in pods from plants grown under control (CT), water deficit (WD), heat stress (HS), or WD+HS. All transcripts shown are significant at P < 0.05 (negative binomial Wald test followed by Benjamini–Hochberg correction). **(B)** Stomatal aperture of pods from plants subjected to CT, HS, WD, or WD+HS, 60 min following application of 7.5 or 0 μM ABA. All experiments were conducted with 3 biological repeats, each with 10 plants as technical repeats. Twenty microscopic fields from all parts of pods were measured for each plant. **(C)** Inner temperature of pods from plants subjected to CT, WD, HS, or WD+HS, coated or uncoated with a thin layer of petroleum jelly (PTJ) for 3 hours. Results are shown as box-and-whisker plots with borders corresponding to the 25^th^ and 75^th^ percentiles of the data. Different letters denote significance at P < 0.05 (ANOVA followed by a Tukey’s post hoc test). Abbreviations: ABA, abscisic acid; CT, control; HS, heat stress; PTJ, petroleum jelly, WD, water deficit.

As shown in Figure 4B, external application of low levels of ABA caused stomatal closure in pods from CT plants, while stomata of pods from WD stressed plants appeared unresponsive to this treatment and remained closed. In contrast, and in agreement with our previous findings with flowers (Sinha et al., 2022), stomata on pods from plants subjected to HS and WD+HS remained partially open even after external ABA application, maintaining an aperture that did not differ from that of pods of CT plants (Figure 4B). As insensitivity to ABA could also result from suppressed expression of ABA receptors, or key ABA signaling components, in pods during WD+HS, we checked the expression of several transcripts involved in ABA sensing and responses in our RNA-Seq dataset. As shown in Supplemental Figure S2, many transcripts involved in ABA perception/ signaling [*i*.*e*., pyrabactin resistance (PYR)/PYR-like (PYL), protein phosphatase 2C (PP2C), and SNF1-related protein kinase 2 (SnRK2)] are expressed in pods during a combination of WD+HS. In addition, many ABA-response transcripts [*i*.*e*., ABA Insensitive 5 (ABI5), dehydration-responsive element binding (DREB), and responsive to dehydration 22 (RD22)] are also expressed in pods during WD+HS (Supplemental Figure S2).

To determine whether the opening of stomata on pods under conditions of WD+HS indeed contributes to the lowering of internal pod temperature (Figures 1 and 2), we applied a thin layer of petroleum jelly (PTJ; Sinha et al., 2022) to seal stomata on pods of plants subjected to CT, WD, HS and WD+HS, and measured internal pod temperatures. As shown in Figure 4C, PTJ application to pods from CT, HS, and WD+HS plants resulted in elevated internal pod temperature (highest elevation in pods from plants subjected to HS and WD+HS), while PTJ application to pods from WD plants did not.

### Seed size of soybean plants subjected to WD+HS is larger than that of plant subjected to HS

To investigate whether differential transpiration of pods (Figures 1-4) provided some form of protection to seed development under conditions of WD+HS, we measured the total flower, pod and seed numbers per plant, and the mass per seed of soybean plants, growing under conditions of CT, WD, HS and WD+HS. As shown in Figure 5A-5C, compared to plants subjected to CT, WD, or HS, plants subjected to WD+HS produced fewer flowers, pods, and seeds per plant. In contrast, the average mass of seeds from plants subjected to WD+HS was greater compared to that of plants subjected to HS alone (Figure 5D). In addition, when the different seeds developing within pods from the different plants were scored for size (large, medium, and small; Figure 5E-G), it was found that small, developmentally suppressed or potentially aborted seeds, were only found in pods from plants subjected to HS.

**Figure 5.**
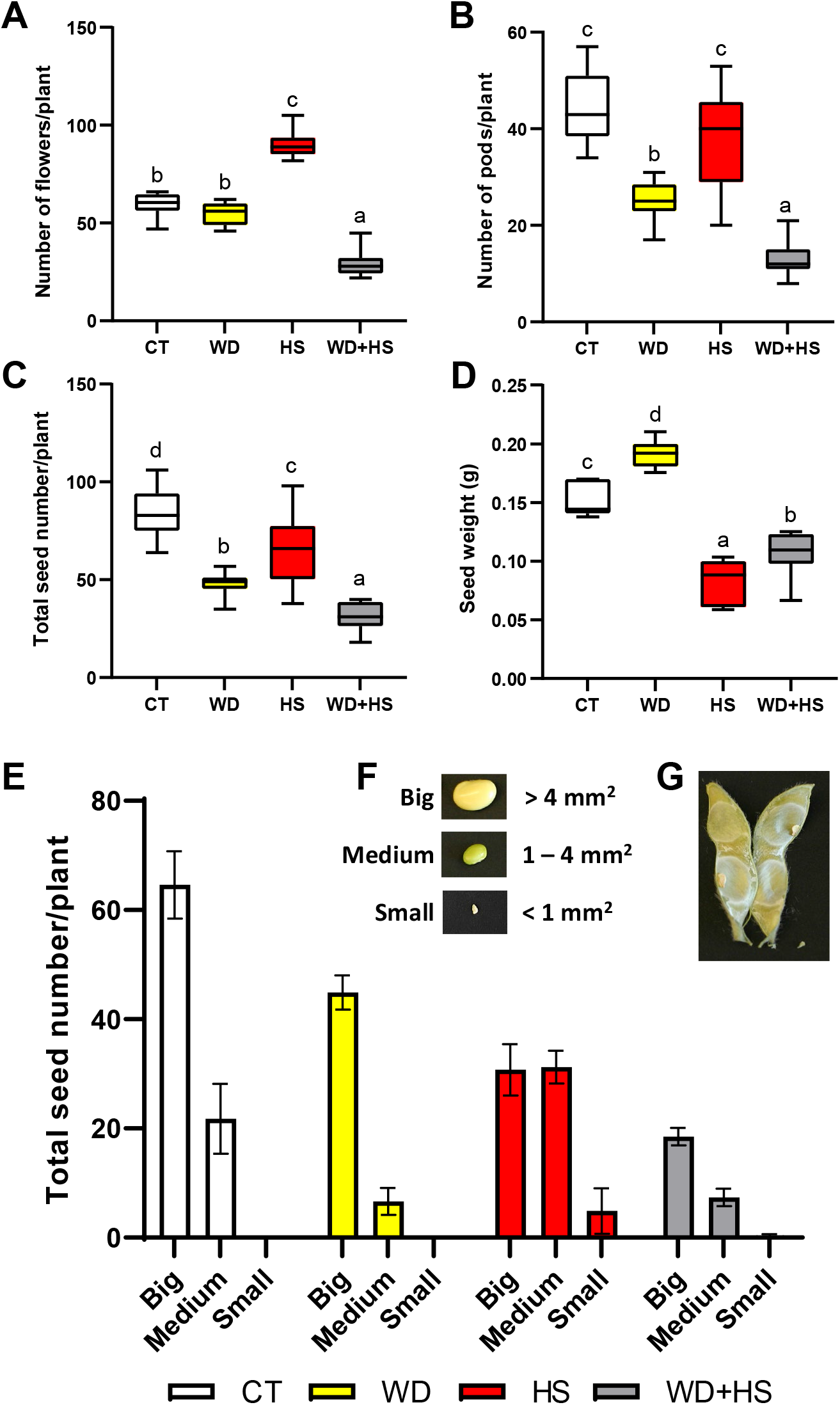
Number or flowers, pods, and seeds per plant and seed mass of plants subjected to a combination of water deficit and heat stress conditions. Total number of flowers **(A)**, pods **(B)**, and seeds **(C)** per plant, in plants grown under conditions of CT, WD, HS, or WD+HS. **(D)** Average **s**eed mass (weight) of seeds from plants subjected to CT, WD, HS, or WD+HS. **(E)** Seed size distribution in pods from plants subjected to the different stress treatments. **(F)** Representative pictures of seeds from the different sizes scored in (E). **(G)** Representative picture of a pod with small seeds obtained from plants subjected to HS. All experiments were conducted with 3 biological repeats, each with 14 plants as technical repeats. Results are shown as bar graphs or box-and-whisker plots with borders corresponding to the 25^th^ and 75^th^ percentiles of the data. Different letters denote significance at P < 0.05 (ANOVA followed by a Tukey’s post hoc test). Abbreviations: CT, control; HS, heat stress; WD, water deficit.

## DISCUSSION

Pod development and seed maturation play a key role in determining the overall yield of soybean plants and other legumes. Similar to flower differentiation and plant fertilization, pod development and seed maturation are negatively impacted by HS (Siebers et al., 2015; Sehgal et al., 2018; Djanaguiraman et al., 2019, 2020). Thus, processes such as embryo development, seed number (a potential result of embryo abortion/death), seed filling, and seed maturation are negatively affected by heat stress, resulting in reduced yield. We previously reported that closed flowers of soybean (a cleistogamous/pseudocleistogamous plant, in which fertilization occurs in closed flowers; Takahashi et al., 2001; Khan et al., 2008) use a strategy of differential transpiration to limit overheating of reproductive tissues (Sinha et al., 2022). Here we report that pods of soybean plants grown under a combination of WD+HS use a similar strategy of differential transpiration to buffer pod internal temperature (Figures 1, 2, and 4). We further show that like flowers (sepals; Sinha et al., 2022), the stomata of pods from plants grown under a combination of WD+HS are less sensitive to external application of ABA than those of CT plants, suggesting that an active process of ABA degradation could drive differential transpiration between leaves and pods under these growth conditions (Figures 1, 2 and 4; Supplemental Figure S2). In addition, compared to plants grown under CT or WD conditions, the stomatal density and index of pods developing on plants under conditions of HS or WD+HS was high (Figure 1), further suggesting that stomatal function is important for this process to occur. A comparison of the expression pattern of transcripts involved in ABA biosynthesis and degradation between pods and flowers grown under the different growth conditions (Figure 4A) revealed however that in contrast to flowers, pods do not have an elevated expression of several different ABA biosynthesis genes (especially noticeable during WD), suggesting that the levels of ABA in pods could be, at least in part, a result of transport from roots or other plant tissues (some transcripts of ABA degrading enzymes were nevertheless upregulated in pods or flowers subjected to WD+HS; Figure 4A; Sinha et al., 2022).

Interestingly, the expression of many different WD-or HS-response transcripts was lower in pods compared to flowers or leaves, in response to WD or HS (Figure 3). In contrast, the response of pods to a combination of WD+HS was intense and included over 11,000 transcripts (Figure 3). The lower number of WD-response transcripts expressed in pods, compared to flowers and leaves, could suggest that due to their different anatomy, as well as being a prime sink tissue of the plant, pods might experience less water stress compared to leaves and flowers, under conditions of WD. In contrast to WD, pods displayed a stronger transcriptomic response to HS (albeit still containing a lower number of transcripts compared to flowers and leaves; Figure 3C), suggesting that pods experience HS in a relatively similar manner as other plant tissues. Interestingly, when WD was combined with HS (WD+HS) the transcriptomic response of all three plant tissues was very high with 1,000s of transcripts altered in their expression (Figure 3). This finding suggests that although pods were not as responsive to WD as leaves and flowers, when WD was combined with HS the stress level of pods was much higher, potentially do to the combined increase in internal temperature and water potential (Figure 1). Comparing the transcriptomic responses of all three tissues to the three different stress conditions, further revealed that each tissue has a distinct response to each stress condition and that the responses of each tissue differ from each other (Figure 3). This finding is very important since it suggests that developing crops with augmented tolerance to climate change may involve altering the responses of each tissue differently. Thus, molecular strategies that will alter the transcriptome or metabolome of the whole plant may work for specific tissues, but not others. Because enhancing yield requires improving leaf-associated vegetative growth, flower-associated reproduction and fertilization processes, and pod-associated seed filling and maturation (as well as root- and other tissues-associated processes), in plants growing under stress, specific engineering solutions might be needed for each plant tissue (leaf, flower, pod, and other tissues). Further studies are therefore needed to identify key signaling and acclimation pathways in each of the different plant tissues involved in the response of plants to stress. In addition, tissue- and stress-specific promoters will be needed to alter these pathways in a stress- and/or tissue-specific manner. One potential strategy to improve yield under stress combination, originating from this work, and the work of Sinha et al., (2022), could involve augmenting the differential transpiration of flowers and pods. For this purpose, the number of stomata on flowers and pods, as well as the degradation rate of ABA in these tissues, could be increased in plants subjected to HS or WD+HS, to improve reproductive tissue cooling [*e*.*g*., by constitutive, or stress-induced, augmented expression of the ABA degradation enzyme CYP707A specifically in pods and flowers, using pod- and flower-specific promoters]. In addition to WD+HS combination, this strategy could also work for other stress combinations that result in stomatal closure under HS conditions (*e*.*g*., combinations of pathogen infection, mechanical injury, high CO_2_, or air pollution, such as ozone, that cause stomatal closure, with HS; Mittler and Zandalinas, 2022). In addition to future studies in soybean, the impact of WD+HS combination on seed filling and maturation, and the potential of differential transpiration to protect these processes, should be studied in other major crops. Conditions of HS and WD+HS are expected to increase in frequency and intensity in the coming years and protecting crop reproductive processes from these conditions should be a prime directive of breeders, biotech industry and academia (Mazdiyasni and AghaKouchak, 2015; Alizadeh et al., 2020; Rivero et al., 2021; Zandalinas et al., 2021; Mittler and Zandalinas, 2022).

An additional interesting finding of this study is that seed mass of plants subjected to WD+HS is larger than that of plants subjected to HS (Figure 5D). This phenotype could be a result of diverting more resources to seed production under conditions of WD+HS (compared to HS), due to differential transpiration (Figures 1-4) and the presence of a lower number of flowers and pods per plant (that do not occur during HS in well-watered plants; Figure 5A, 5B). These could bring more resources such as nutrients from roots or leaves to the developing pods. In addition, compared to plants subjected to HS, no evidence of suppressed seed development, seed filling, or seed abortion, in the form of small seeds, was found in pods from plants subjected to WD+HS (Figure 5E). This finding could also suggest that seeds developing on plants subjected to WD+HS are better protected and/or better nourished due to the process of differential transpiration and/or the presence of fewer pods on each plant (Figure 5). Further studies are needed to determine the potential of these small seeds to germinate and the overall effects of WD+HS on seed filling, abortion, and development under conditions of WD+HS.

Taken together, our findings reveal that, compared to leaves, pods display differential transpiration during conditions of WD+HS. This strategy reduces pod temperature by about 4 °C and likely alleviates high temperature effects on processes such as differentiation, seed filling and maturation that occur within pods, thus limiting detrimental impacts on yield.

## MATERIALS AND METHODS

### Soybean growth and stress treatments

Soybean (*Glycine max*, cv *Magellan*) seeds were inoculated with *Bradyrhizobium japonicum* inoculum (N-DURE, Verdesian Life Sciences, NC, USA) and germinated in Promix BX (Premier Tech Horticulture; PA, USA), for a week in a growth chamber (BDR16, Conviron; Canada) under short day growth condition (12-h light/12-h dark), at 28/24 °C day/night temperature and 500 μmol photons m^-2^ s^-1^. The temperature of the chambers was ramped from 24 to 28 °C between 6.00-8.00 AM and decreased to 24 °C from 16.00-20.00 PM. Seedlings were transplanted into pots containing 1 kg mixture of Promix BX and perlite (Miracle-Gro® Perlite, Miracle-Gro, Marysville, OH, USA) mixed in ratio of 10:1 and soaked in 1 l of water-fertilizer (Zack’s classic blossom booster 10-30-20; JR peters Inc., USA) mix (Cohen et al., 2021a, Sinha et al., 2022). Plants were then grown under 28/24 °C day/night temperatures and 1000 μmol photons m^-2^ s^-1^ light intensity (12-h light/12-h dark photoperiod) for the next 16-18 days (until start of first open flower, R1 developmental stage, Fehr et al., 1971) while irrigating twice a week with fertilizer (Cohen et al., 2021a, Sinha et al., 2022). At R1, plants were randomly divided into control (CT), and 3 stress treatments of water-deficit (WD), heat stress (HS), and a combination of water-deficit and heat stress (WD+HS) in four identical BDR16 growth chambers placed side-by-side in the same room (Sinha et al., 2022; 15 plants per chamber). The relative humidity of chambers was maintained at about 50-60% in all chambers. The plants under WD and WD+HS treatments were irrigated with only 30% of the water available for transpiration as described previously (Cohen et al., 2021a, Sinha et al., 2022), while plants in the CT and HS treatments were irrigated with 100% of the water available for transpiration. For HS and WD+HS treatments, the temperatures in the chambers were maintained at 38 °C day and 28 °C night temperature by gradually increasing the temperature between 6.00-8.00 AM and decreasing it between 16.00-20.00 PM, to achieve the indicated day and night temperatures.

### Temperature, gas exchange and water potential

Pod internal temperature was measured between 11.30 AM-12:30 PM using a microthermocouple sensor (Physitemp instruments LLC; Clifton, NJ, USA), attached to a Multi-Channel Thermocouple Temperature Data Logger (TCTemp X-Series, ThermoWorks LogMaster; UT, USA; Sinha et al., 2022). The hypodermal needle microprobe (Physitemp instruments LLC; Clifton, NJ, USA) of the microthermocouple sensor was inserted 3-4 mm into the soybean pods and the temperature of the internal pod cavity and developing seeds was averaged for each pod. Transpiration and stomatal conductance of pods and leaves were measured using a LICOR Portable Photosynthesis System (LI-6800, LICOR, Lincoln, NE, USA) between 12.00-1.00 PM as described previously for soybean flowers (Sinha et al., 2022). Water potential of pods (cut open into half) from the different stress treatments was measured using a Dewpoint Potentiometer (WP4C, METER Group, Inc. WA, USA) as described previously (Cohen et al., 2021a, Sinha et al., 2022).

### Stomatal measurements and sealing of stomata

Abscisic acid (ABA, 7.5 μM, Sigma-Aldrich, St. Louis, MO, USA) was sprayed on pods of soybean plants growing under the different stress conditions, as previously described for soybean flowers (Sinha et al., 2022), while leaves were shielded with a plastic layer. For control, pods from plants grown under the different conditions were sprayed with water (Sinha et al., 2022). Plants were then returned to the chambers and 60 min post ABA application the pod surface was quickly covered with thin layer of transparent nail polish (Sally Hansen topcoat, Sally Hansen, NY, USA). Once, the nail polish layer was dry, it was removed from pods and placed on a microscope slide, covered with another microscopic slide and stomata images were recorded using an EVOS XL microscope (Invitrogen by Thermo Fisher Scientific, CA, USA) as described previously (Devireddy et al., 2020; Zandalinas et al., 2020, Sinha et al., 2022, Xie et al., 2022). The width and length of stomatal aperture were measured using ImageJ (https://imagej.nih.gov/ij) and stomatal aperture was calculated as ratio of stomatal pore width to stomatal pore length (Sinha et al., 2022). Number of epidermal and pavement cells per microscopic field of view were counted using ImageJ to calculate stomatal density and stomatal pore index as previously described (Sinha et al., 2022). To inhibit transpirational cooling, the entire surface of pods from soybean plants grown under the different growth conditions were sealed by gently applying a thin layer of petroleum jelly (Vaseline®; Sigma-Aldrich, St. Louis, MO, USA) using Q-tips (Sinha et al., 2022). Plants were then returned to the chambers and pod temperatures were measured 3 hrs post petroleum jelly application using microthermocouple as described above.

### RNA isolation, sequencing, and data analysis

Pods of soybean plants grown under the different conditions were collected between 11.30 AM-12:30 PM. Samples were flash frozen in liquid nitrogen. For each biological repeat, pods from 8-10 different plants were pooled and RNA was isolated using RNAeasy plant mini kit (Qiagen, Germantown, MD, USA). RNA libraries were prepared using standard Illumina protocols and RNA sequencing was performed using NovaSeq 6000 PE150 by Novogene co. Ltd (https://en.novogene.com/; Sacramento, CA, USA). Read quality control was performed using Trim Galore v0.6.4 (https://www.bioinformatics.babraham.ac.uk/projects/trim_galore/) & FastQC v0.11.9 (https://www.bioinformatics.babraham.ac.uk/projects/fastqc/). The RNA-seq reads were aligned to the reference genome for Soybean - Glycine max v2.1 (downloaded from ftp://ftp.ensemblgenomes.org/pub/plants/release-51/fasta/glycine_max/dna/), using Hisat2 short read aligner (Kim et al., 2019). Intermediate file processing of sam to sorted bam conversion was carried out using samtools v1.9 (Danecek et al., 2021). Transcript abundance in levels expressed as FPKM was generated using the Cufflinks tool from the Tuxedo suite (Trapnell et al., 2012), guided by genome annotation files downloaded from the same source. Differential gene expression analysis was performed using Cuffdiff tool (Trapnell et al., 2013), also from the same Tuxedo suite. Differentially expressed transcripts were defined as those that had a fold change with an adjusted P < 0.05 (negative binomial Wald test followed by Benjamini–Hochberg correction). Functional annotation and quantification of overrepresented GO terms (P value < 0.05) were conducted using g:profiler (Raudvere et al., 2019). Venn diagrams were created in VENNY 2.1 (BioinfoGP, CNB-CSIC). Venn diagram overlaps were subjected to hypergeometric testing using the R package phyper (Zandalinas et al., 2020a). Heatmaps were generated in Morpheus (https://software.broadinstitute.org/morpheus). The transcriptome response of soybean leaves, flowers, and pods, to CT, WD, HS, and WD+HS conditions was studied using the same growth chambers, experimental conditions, and RNA-seq analysis and quantification protocols described above and in Sinha et al., (2022).

### Quantification of yield components

Pods were collected from 13-14 soybean plants growing under the different stress and control condition 42 days after initiation of the stresses. Flower and pod numbers per plant were measured as described in (Sinha et al., 2022). To count the number of seeds with different sizes, the area of each seed was measured using ImageJ (imagej.nih.gov/ij) and the number of seeds with surface area more than 4 mm^2^, between 1-4 mm^2^, or lower than 1 mm^2^ were classified as big, medium, and small (developmentally arrested or potentially aborted) seeds, respectively. Seed mass was measured by weighing each individual seed from each plant using an analytical scale.

### Statistical Analysis

All experiments were repeated 3 times (biological repeat), each time with at least 15 plants as technical repeats. Results are presented as box-and-whisker plots with borders corresponding to the 25^th^ and 75^th^ percentiles of the data. Statistical analysis was performed using one-way ANOVA followed by Tukey’s post hoc test (P < 0.05) in GraphPad (Sinha et al., 2022). Different letters denote statistical significance at P < 0.05.

## Data Availability

The analyzed transcript abundance and differentially expressed transcripts can be accessed interactively via Differential Expression tool in SoyKB; https://soykb.org/DiffExp/diffExp.php; Joshi et al., 2012, 2014), a comprehensive web resource for soybean. It provides a set of visualization and analytical tools such as differential expression analysis and gene card pages and provides data in the form of tabs for Gene lists, Venn diagram, Volcano plot, Function Analysis, Pathway Analysis and Gene modules. RNA-seq sequence data from this article can be found in the Gene Expression Omnibus (GEO) database under the accession number: GSE213479.

## Accession Numbers

Sequence data from this article can be found in the GenBank/EMBL data libraries under accession numbers: *ABA2 – Glyma*.*11G151700 / Glyma*.*11G151400/NM_104113*.*5; ABA1 – Glyma*.*09G000600/ Glyma*.*11G055700/ Glyma*.*17G174500/NM_180954*.*3, CYP707A – Glyma*.*16G109300/Glyma*.*02G132200/ Glyma*.*07G212700/Glyma*.*17G242200/NM_118043*.*2, NCED3 – Glyma*.*15G250100/NM_112304*.*3, NCED4 – Glyma*.*01G154900/NM_118036*.*3, NCED5 – Glyma*.*05G140900/Glyma*.*08G096200/NM_102749*.*3, AAO3 – Glyma*.*02G272200/NM_128273*.*3*.

## Supplemental Data

**Supplemental Figure S1**. Venn diagrams showing the overlap in expression of selected transcription factor families in pods, flowers, and leaves from plants grown under CT, WD, HS and WD+HS conditions.

**Supplemental Figure S2**. Expression of transcripts involved in ABA perception and signaling in pods from plants subjected to WD, HS, or WD+HS.

**Supplemental Dataset S1**. Transcripts upregulated in soybean pod subjected to WD stress (Figure 3A).

**Supplemental Dataset S2**. Transcripts downregulated in soybean pod subjected to WD stress (Figure 3A).

**Supplemental Dataset S3**. Transcripts upregulated in soybean pod subjected to HS (Figure 3A).

**Supplemental Dataset S4**. Transcripts downregulated in soybean pod subjected to HS (Figure 3A).

**Supplemental Dataset S5**. Transcripts upregulated in soybean pod subjected to WD+HS (Figure 3A).

**Supplemental Dataset S6**. Transcripts downregulated in soybean pod subjected to WD+HS (Figure 3A).

**Supplemental Dataset S7**. GO enrichment categories of transcripts unique to WD+HS in pod (Figure 3B).

**Supplemental Dataset S8**. Transcripts exclusively differentially expressed in soybean pod subjected to WD (Figure 3D).

**Supplemental Dataset S9**. Transcripts exclusively differentially expressed in soybean flower subjected to WD (Figure 3D).

**Supplemental Dataset S10**. Transcripts exclusively differentially expressed in soybean leaf subjected to WD (Figure 3D).

**Supplemental Dataset S11**. Transcripts exclusively differentially expressed in soybean pod subjected to HS (Figure 3D).

**Supplemental Dataset S12**. Transcripts exclusively differentially expressed in soybean flower subjected to HS (Figure 3D).

**Supplemental Dataset S13**. Transcripts exclusively differentially expressed in soybean leaf subjected to HS (Figure 3D).

**Supplemental Dataset S14**. Transcripts exclusively differentially expressed in soybean pod subjected to WD+HS (Figure 3D).

**Supplemental Dataset S15**. Transcripts exclusively differentially expressed in soybean flower subjected to WD+HS (Figure 3D).

**Supplemental Dataset S16**. Transcripts exclusively differentially expressed in soybean leaf subjected to WD+HS (Figure 3D).

**Supplemental Dataset S17**. GO enrichment categories of transcripts unique to CT in pod, flower, and leaf.

**Supplemental Dataset S18**. GO enrichment categories of transcripts unique to WD in pod, flower, and leaf.

**Supplemental Dataset S19**. GO enrichment categories of transcripts unique to HS in pod, flower, and leaf.

**Supplemental Dataset S20**. GO enrichment categories of transcripts unique to WD+HS in pod, flower, and leaf.

**Supplemental Dataset S21**. Differential expression of heat shock factor (HSF) transcripts in soybean pod, flower and leaf subjected to WD, HS and WD+HS (Figure 3E).

**Supplemental Dataset S22**. Differential expression of AP2/ERF/RAV transcription factor transcripts in soybean pod, flower and leaf subjected to WD, HS and WD+HS (Figure 3E).

**Supplemental Dataset S23**. Differential expression of MYB transcription factor transcripts in soybean pod, flower and leaf subjected to WD, HS and WD+HS (Figure 3E).

**Supplemental Dataset S24**. Differential expression of NAC transcription factor transcripts in soybean pod, flower and leaf subjected to WD, HS and WD+HS (Figure 3E).

**Supplemental Dataset S25**. Differential expression of WRKY transcription factor transcripts in soybean pod, flower and leaf subjected to WD, HS and WD+HS (Figure 3E).

**Supplemental Dataset S26**. Differential expression of bHLH transcription factor transcripts in soybean pod, flower and leaf subjected to WD, HS and WD+HS (Figure 3E).

**Supplemental Dataset S27**. Differential expression of ARF transcription factor transcripts in soybean pod, flower and leaf subjected to WD, HS and WD+HS (Figure 3E).

**Supplemental Dataset S28**. Differential expression of CAMTA transcription factor transcripts in soybean pod, flower and leaf subjected to WD, HS and WD+HS (Figure 3E).

---

**Supplementary Fig S1.**
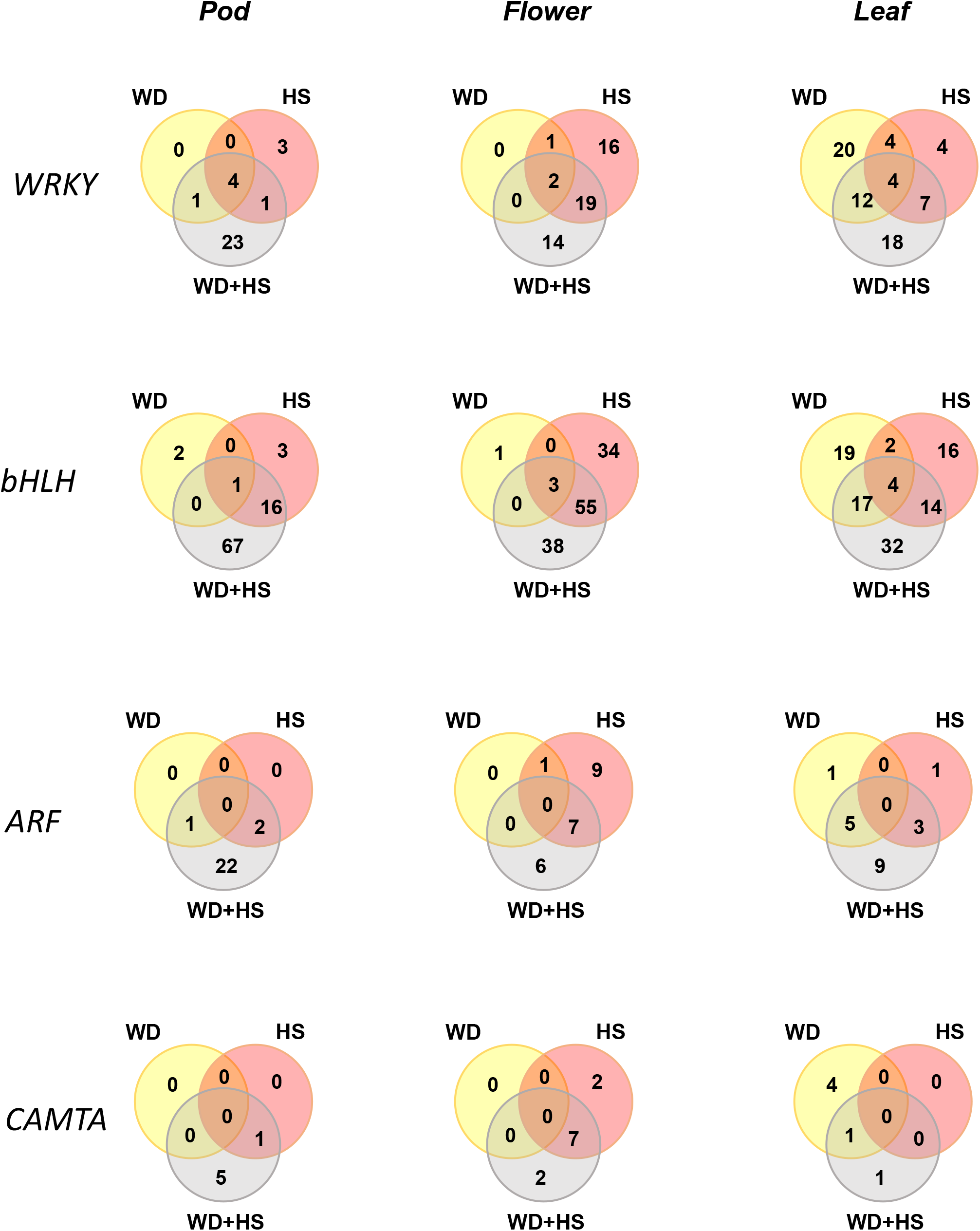
Comparison of transcription factor gene expression between WD, HS and WD+HS in pod, flower and leaf (in support of Figure 3). Venn diagrams showing the overlap in expression of selected transcription factor (TF) families in pods, flowers, and leaves from plants grown under CT, WD, HS and WD+HS conditions. Abbreviations: ARF, Auxin response factors; bHLH, basic helix-loop-helix; CAMTA, calmodulin binding transcription activator; CT, control; HS, heat stress; WD, water deficit.

**Supplementary Fig S2.**
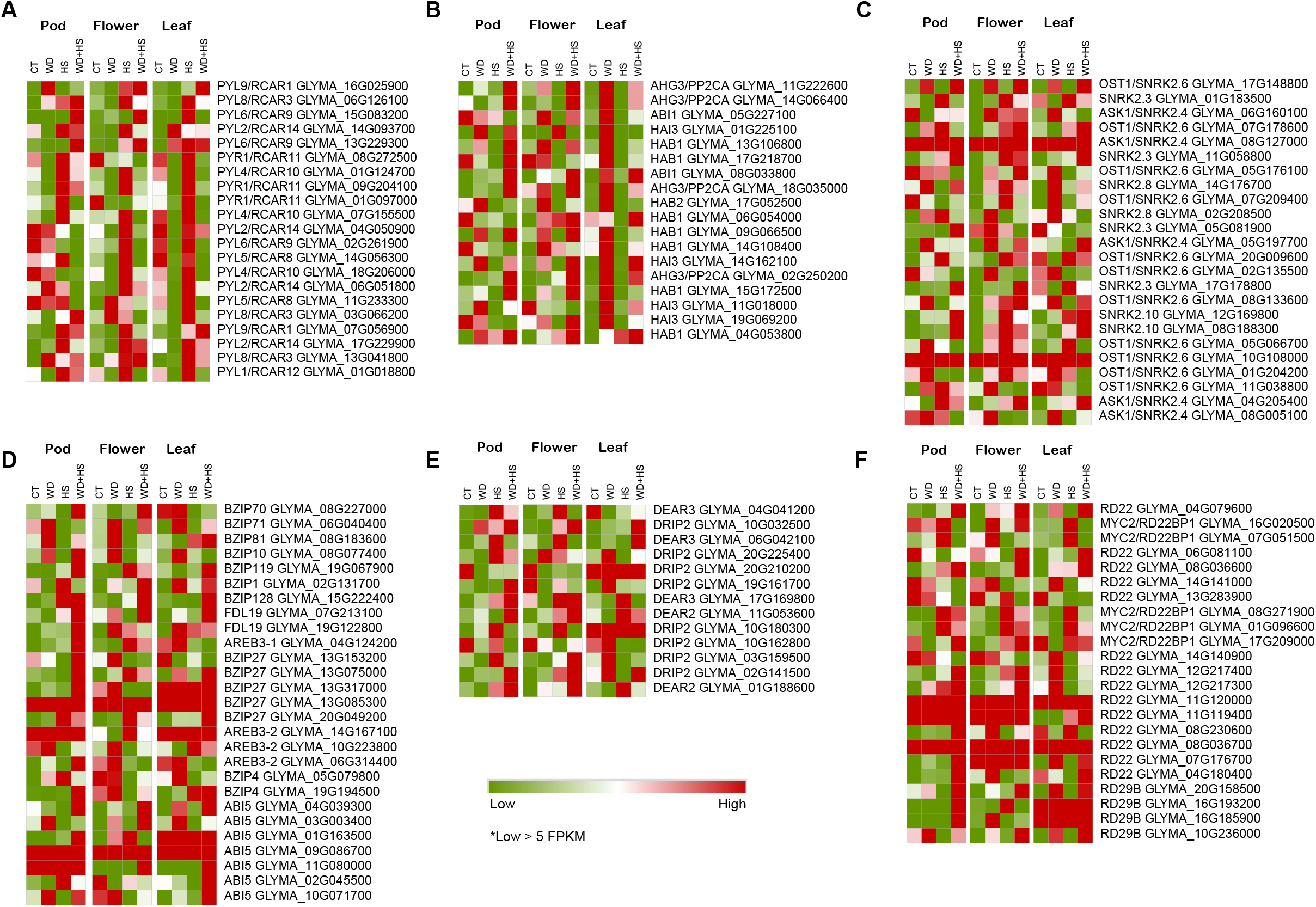
Expression of transcripts involved in ABA perception and signaling in leaves, flower and pods from plants subjected to WD, HS, and WD+HS (in support of Figure 4). Heat maps showing the expression level of transcripts involved in ABA perception: ABA receptor PYL/PYR **(A)**; PP2C **(B)**; SNRK2 **(C)**, and ABA signaling: ABI5 **(D)**; DREB **(E)**; RD22 **(F)**, in pod, flower and leaf tissue from plants subjected to WD, HS, and WD+HS. Expression level is in FPKM was standardized across the entire experiment as described in methods. Abbreviations: CT, control; HS, heat stress; WD, water deficit; PYR, pyrabactin resistance; PYL, PYR-like; PP2C, protein phosphatase 2C; SnRK2, SNF1-related protein kinase 2; ABI5, ABA Insensitive 5; DREB, dehydration-responsive element binding; RD22, responsive to dehydration 22; FPKM, fragments per kilobase of exon per million mapped fragments.

